# Gene-level quantitative trait mapping in an expanded *C. elegans* multiparent experimental evolution panel

**DOI:** 10.1101/589432

**Authors:** Luke M. Noble, Matthew V. Rockman, Henrique Teotónio

**Author notes:** Corresponding author: Institut de Biologie de l’École Normale Supérieure (IBENS), Inserm U1024, CNRS UMR 8197, PSL Research University, F-75005 Paris, France.

## Abstract

The *Caenorhabditis elegans* multiparental experimental evolution (CeMEE) panel is a collection of genome-sequenced, cryopreserved recombinant inbred lines useful for mapping the genetic basis and evolution of quantitative traits. We have expanded the resource with new lines and new populations, and here report updated additive and epistatic mapping simulations and the genetic and haplotypic composition of CeMEE version 2. Additive QTL explaining 3% of trait variance are detected with >80% power, and the median detection interval is around the length of a single gene on the highly recombinant chromosome arms. Although CeMEE populations are derived from a long-term evolution experiment, genetic structure is dominated by variation present in the ancestral population and is not obviously associated with phenotypic differentiation. *C. elegans* provides exceptional experimental advantages for the study of phenotypic evolution.

## Introduction

Multiparent recombinant inbred line panels (MPP) are key components of the tool-kit employed by quantitative geneticists to find quantitative trait loci (QTL) estimate trait heritability and predict phenotype values, and to study the genetics of phenotypic evolution (Lynch and Walsh 1998; Barton and Keightley 2002). MPP allow experimenters to sample more variation than that present in biparental recombinant inbred lines (RILs) and, subject to effective recombination – itself dependent on breeding mode and population sizes, and meitoic features such as crossover interference and karotype – greatly increase QTL mapping resolution (Valdar *et al.* 2006; Rockman 2008). When compared to genome-wide association studies (GWAS) usually performed in natural populations of often unclear pedigree and evolutionary history, including human populations, MPP allows one to measure individual breeding values and to control for potentially confounding environmental covariates and population structure due to drift, mating or reproductive system, or natural selection.

We recently introduced the *Caenorhabditis elegans* Multiparental Experimental Evolution (CeMEE) panel as the first MPP for this model organism, and described the polygenic epistatic genetic architecture of fertility and body size (Noble *et al.* 2017). The CeMEE panel is derived from an intercross of 16 wild isolate founders and between 140 and 240 generations of subsequent experimental evolution under variable sex ratios and breeding mode (self-fertilization and outcrossing), at census population sizes of 10^4^ and effective population sizes of around 10^3^ (Chelo et *al.* 2013), in a defined laboratory environment that is fairly stable and unstructured (Teotónio et *al.* 2012). For these reasons, hybrid genomes are a well-recombined mix of the 16 founders and traits can be appropriately assayed for their relation with fitness (Teotónio *et al.* 2017). Now comprising 733 genome-sequenced RILs, the CeMEE panel has good power to map additive QTL explaining >3% of trait variance, at a mapping resolution of 5-20Kb depending chromosomal region, values comparable to those of the Drosophila Synthethic Population Resource (DSPR), arguably the best metazoan MPP so far (King *et al.* 2012a, b). Divergence during experimental evolution has created population structure across the CeMEE when viewed as a whole (Noble *et al.* 2017), however the extent of differentiation is surprisingly weak considering the now more than 500 generations that in sum separate the population replicates from which CeMEE RILs were derived.

## Methods

### Experimental evolution and recombinant inbred lines

Building on CeMEE version 1 (Noble *et al.* 2017), we sequenced (or resequenced to greater depth) an additional 455 recombinant inbred lines (RILs). Of these, 169 sequenced lines came from three new populations (control androdioecious CA[1-3]100, where [1-3] designates the population replicate), which were evolved from CA[1-3]50 populations for a further 50 generations under our standard experimental evolution conditions (Figure 1). Other RILs come from the already reported populations: A6140, a lab domesticated population derived from the A0 hybrid population (itself derived by parallel intercrosses among the 16 wild isolates) by 140 generations of experimental evolution (Teotónio *et al.* 2012); and GA[1,2,4], GT[1,2] and GM[1,3] populations, for androdioecious, trioecious and monoecious populations evolved in gradually increasing NaCl concentrations (Theologidis *et al.* 2014). In brief, our standard laboratory environment involved constant census size controlled by seeding each of 10 plates per population with 1000 swimming synchronized L1 larvae, growth at 20° C in the presence of excess *E. coli,* and discrete generations enforced by bleaching of reproductively mature adults at 4 days post seeding (Teotónio *et al.* 2012). Plates was filled with 28mL of Nematode Growth Medium lite (NGM-lite, US Biological) where NaCl concentration was 25 mM (for A6140 and all CA populations), or supplemented with NaCl reaching a maximum concentration of 305 mM for the GA, GT and GM populations from generation 35 to generation 50 (Theologidis *et al.* 2014). As before, the new RILs from CA[1-3] were derived by sampling single hermaphrodites from populations and selfing for 10 generations before preparation of genomic DNA and cryopreservation. The number of RILs derived (sequenced, post-quality control, or currently in cryopreservation but unsequenced) are shown in Table 1.

**Table 1:** Number of recombinant inbred lines derived from experimentally evolved populations in CeMEE v2 and v1 (Noble *et al.* 2017), and the total number of replicated, cryopreserved RILs (CeMEE v2 post-QC plus additional lines with no sequencing data at present). Populations derived from A6140 are formatted as *TMRG* where *T* is evolution treatment (here, Control conditions or Gradual adaptation to a moving optimum), *M* is mating system (Androdioecious, Trioecious or *M*onoecious), *R* is replicate number, and *G* is the number of generations from A6140.

| Source population | v1 | v2 | cryo. |
| --- | --- | --- | --- |
| A6140 | 178 | 248 | 299 |
| CA[1-3]50 | 118 | 143 | 148 |
| CA[1-3]100 | 0 | 128 | 305 |
| GA[1,2,4]50 | 127 | 131 | 152 |
| GT[1,2]50 | 79 | 78 | 78 |
| GM[1,3]50 | 5 | 5 | 5 |
|  | 507 | 733 | 987 |

**Figure 1:**
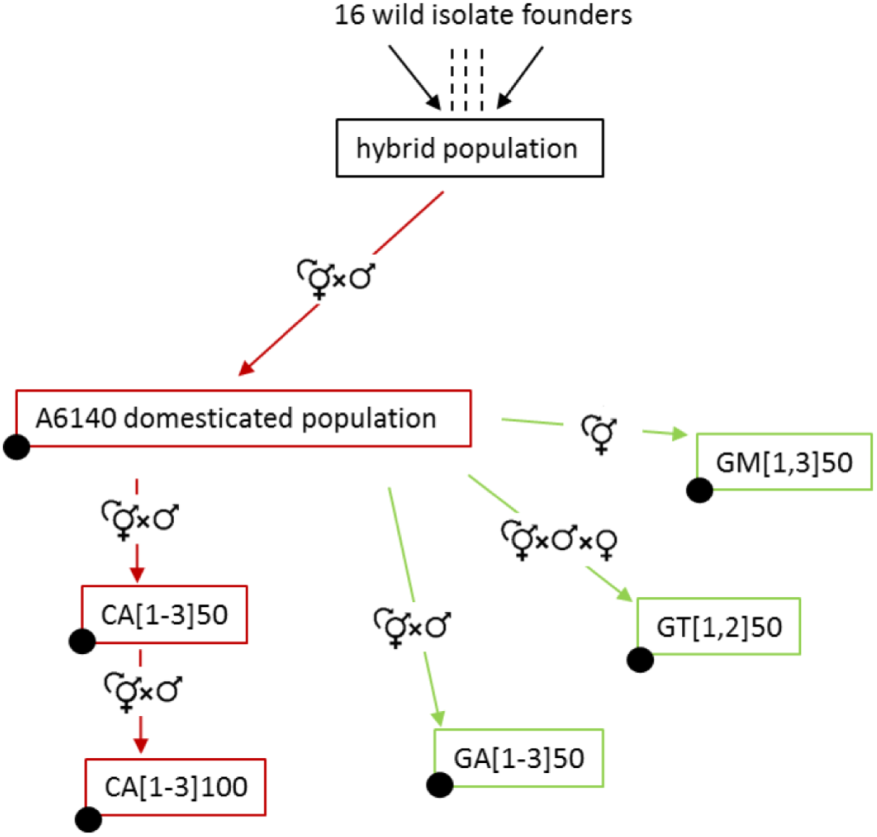
Experimental evolution scheme. Colors indicate specific environmental and reproductive system regimes: black for derivation of the hybrid (A0) population from wild isolates, red for domestication under mixed selfing and outcrossing (A6140), orange for continued evolution in the standard environment (100 generations), green for evolution in changing environments (50 generations). In each regime, samples from replicate populations (numbered in square brackets) are periodically frozen for contemporaneous characterization of ancestral and derived characters. Filled circles indicate populations from which CeMEE RILs were derived.

### Genome sequencing and variant calling

RIL genomic DNA was prepared using the Qiagen Blood and Tissue kit. Library preparation and sequencing was carried out at the Beijing Genome Institute on the Illumina HiSeq X Ten or BGIseq platforms with, respectively, 150 and 100bp paired-end reads to a mean depth of 4.2x and 22x (estimated from per base mapped read depth at the central recombination rate domain of chromosome I).

A set of 388,201 filtered diallelic single nucleotide polymorphisms (SNPs) segregating in the 16 founders was used to impute genotypes by a Hidden Markov Model (HMM; as in (Noble *et al.* 2017)), but given the generally higher sequencing coverage of the new lines we also supplemented imputation with direct variant calling (joint calling with GATK v4.0.2 HaplotypeCaller McKenna *et al.* (2010). We observed high frequencies of heterozygous calls among CA100 RILs (e.g., 93% of 135 lines sequenced to >10x coverage showed both reference and alternative alleles at >20% of segregating sites), particularly for population replicate 3. Allelic proportions were generally low, however (only 11% of these 135 lines had mean proportions >20%), consistent with either DNA contamination or extremely strong selection on widespread residual heterozygosity during amplification for DNA extraction. Given the low levels of heterozygosity seen among other inbred lines sequenced to equivalent depth, including the CA50 ancestors, we consider it unlikely that the CA100 populations have evolved severe inbreeding depression (also supported by unpublished data for fitness-correlated traits). We removed extreme cases as part of our quality control (see below), and define the following genotype sets used as bases of analysis:

1. The full set of 329,976 segregating (i.e., minor allele frequency > 0) diallelic SNPs imputed by HMM, coded [0,1] relative to the WS220 N2 reference genome, with 0.06% of genotype calls of uncertain zygosity (coded > 0 and < 1).
2. As above, but with all genotypes called as heterozygous by GATK coded as 0.5 (1.7% of all genotype calls).

### Data and materials archiving

All raw read data is available from the NCBI SRA under under BioProject PRJNA381203. Sequencing and other metadata for the 16 founders and all RILs, as well as a genome browser and all simulation code used in this paper, are available from lukemn.github.io/cemee. All lines (including founders, CeMEE v2 RILs, and additional RILs that have not yet been sequenced) are cryopreserved in replicated 96-vial plates and are freely available for non-commercial purposes.

### RIL quality control

Our quality control considered homozygosity, sequencing depth, relatedness and haplotype reconstruction likelihood. From 838 cryopreserved lines with any sequence data, identity at segregating sites was thresholded to a maximum of 90% (removing all but 5 lines from monoecious GM populations, and 61 lines from other populations), minimum expected sequencing depth to at least 0.1x (n=2) and, based on segregating sites covered by at least 3 reads, a maximum of 20% where both reference and alternate alleles were seen with mean minor allele frequency >20% (n=24). After sequence and genotype filtering, outlier lines with haplotype reconstruction posterior log likelihoods (see below) below the 0.1 percentile in deviation from the population median for more than three chromosomes (n=11) were also excluded. Haplotype outliers showed no significant population bias; they were always associated with a high minor allele proportion at heterozygous sites, though not always a high frequency of heterozygous sites genome-wide.

### Haplotype reconstruction

We used the RABBIT framework (Zheng *et al.* 2014, 2015) for haplotype reconstruction from founder genotypes (marker set 2, with all heterozygous genotypes set as missing data), as in Noble *et al.* (2017). For each chromosome and population replicate, we estimated maximum likelihood map expansion *(Ra)* by Brent search (Brent 2002) under the fully dependent homolog model *(depModel)* for each line (assuming full homozygosity, and founder and RIL genotype error rates of 0.5%) in Mathematica 11.0.1 (Mathematica 2016). Per marker haplotype posterior probabilities, per line likelihoods, and Viterbi-decoded paths (Viterbi 1967) were then calculated using this value. Outliers (as defined above) were removed and the process was repeated to arrive at final *Ra* values shown in Figure 2.

**Figure 2:**
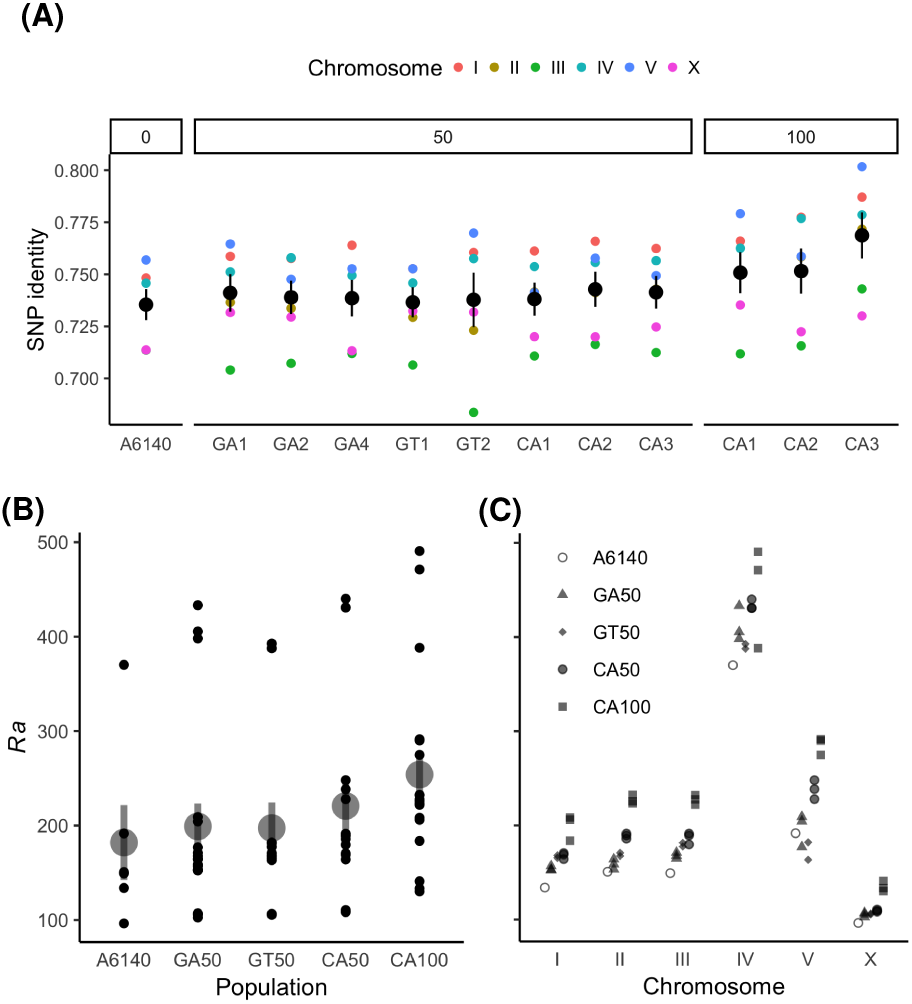
**A.** Genetic relatedness within population replicates, grouped by generation from A6140 (mean and standard error of pairwise identity among lines at segregating sites for each chromosome). **B-C**. Realized map expansion across experimental populations (**B**) and chromosomes (**C**). Each point is a single value for a chromosome of each replicate population, with the mean and standard error overplotted in **B**. Map expansion increases with generation (*p* < 10^−35^ by Poisson linear model), and across all chromosomes, though variably so (*r*^2^ >0.77 for all chromosomes except V (0.58) and the outlying IV (0.37), the latter of which carries a highly recombinant, high frequency haplotype in the right arm piRNA cluster, cf. (Chelo and Teotónio 2013; Noble *et al.* 2017).

**Figure 3:**
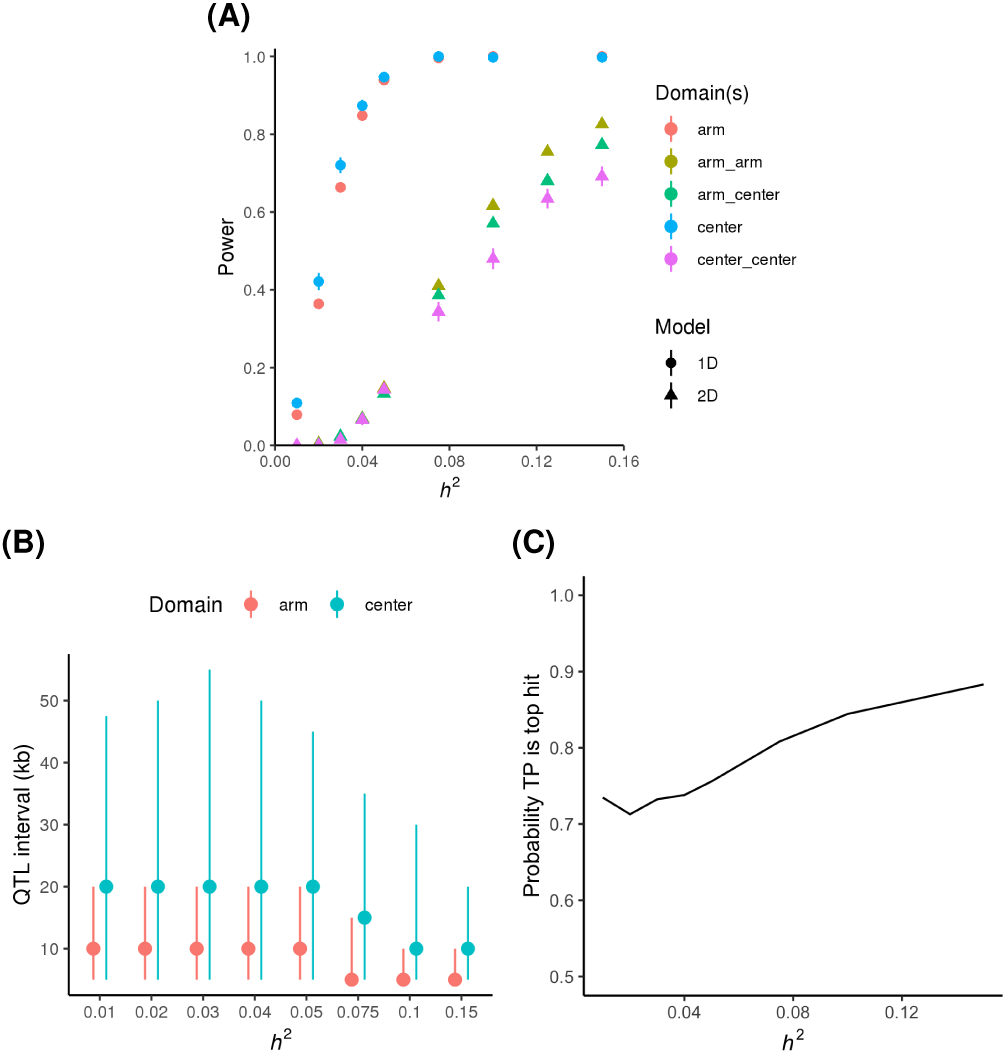
**A**. Detection power for single additive SNPs (1D; random effect set-based model with 5Kb non-overlapping windows, empirical FDR < 1%) or a single interacting marker pair (2D; fixed effects model, 10% FDR based on the effective number of tests). Results (mean and standard error) are split by chromosome recombination rate domain(s) of the simulated site(s). **B**. QTL detection intervals (median and interquartile range) for the additive set test, with the lower bound set by the approximately gene-sized set window. **C.** For additive simulations, the frequency that the lowest p-value genome-wide is that of the window containing the simulated QTL.

**Figure 4:**
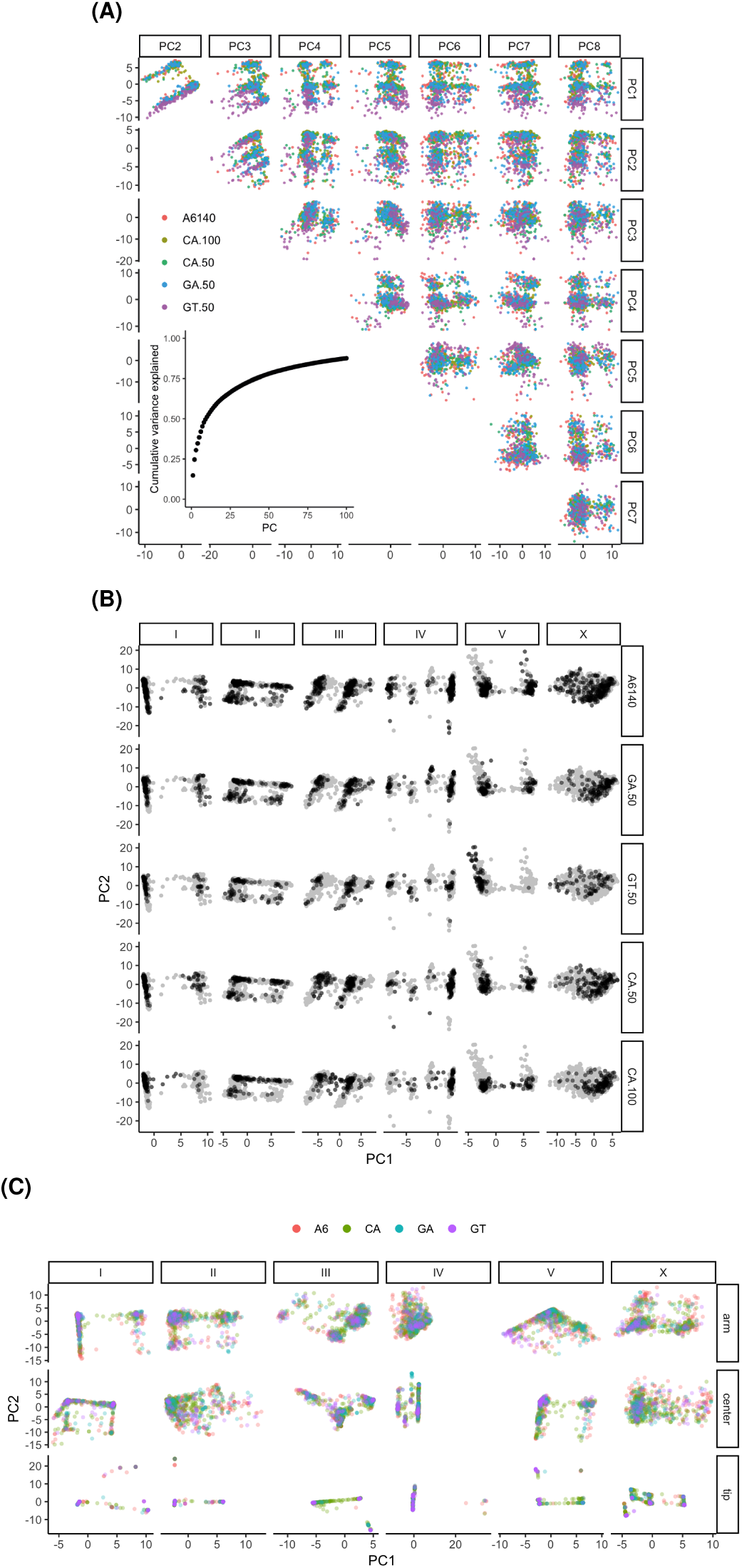
Genetic structure is not dominated by experimental structure. **A**. Biplots for the first eight principal components (PCs) of the additive genetic relatedness matrix (accounting for almost half of the variance), colored by population. The inset shows the cumulative proportion of variance explained by the first 100 PCs. **B**. Populations show relatively consistent structure, with the exception of chromosome V for GT populations. The top PCs for each chromosome are shown for each experimental population (replicates combined), with the full space occupied by all populations in grey. Variance explained ranges from 17% for chromosome X to 57% for IV. **C.** As in **B** but with populations overplotted (CA50 and CA100 pooled), and stratified by recombination rate domain. All values are multiplied by 100.

Haplotype diversity in Figure 5 was summarised from Viterbi paths as entropy across 5Kb windows 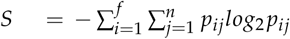, with frequency *p* of haplotype *i* in population *j* summed over *n =* 12 populations and *f* founders (Atwal *et al.* 2007). Haplotype frequencies were evaluated under expectations of pure drift (*f* = 1/16 for equal founder haplotype frequency) or directional selection (frequencies of unique haplotypes from founder genotype data observed in each window). Haplotype differentiation in Figure 5B was based on the variance of the per-population sum of squared deviations from equal founder frequencies, or 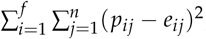, where *p*_*ij*_ is the frequency of haplotype *i* in population *j*, and *e*_*ij*_ is the expected value under drift (1/*f*).

**Figure 5:**
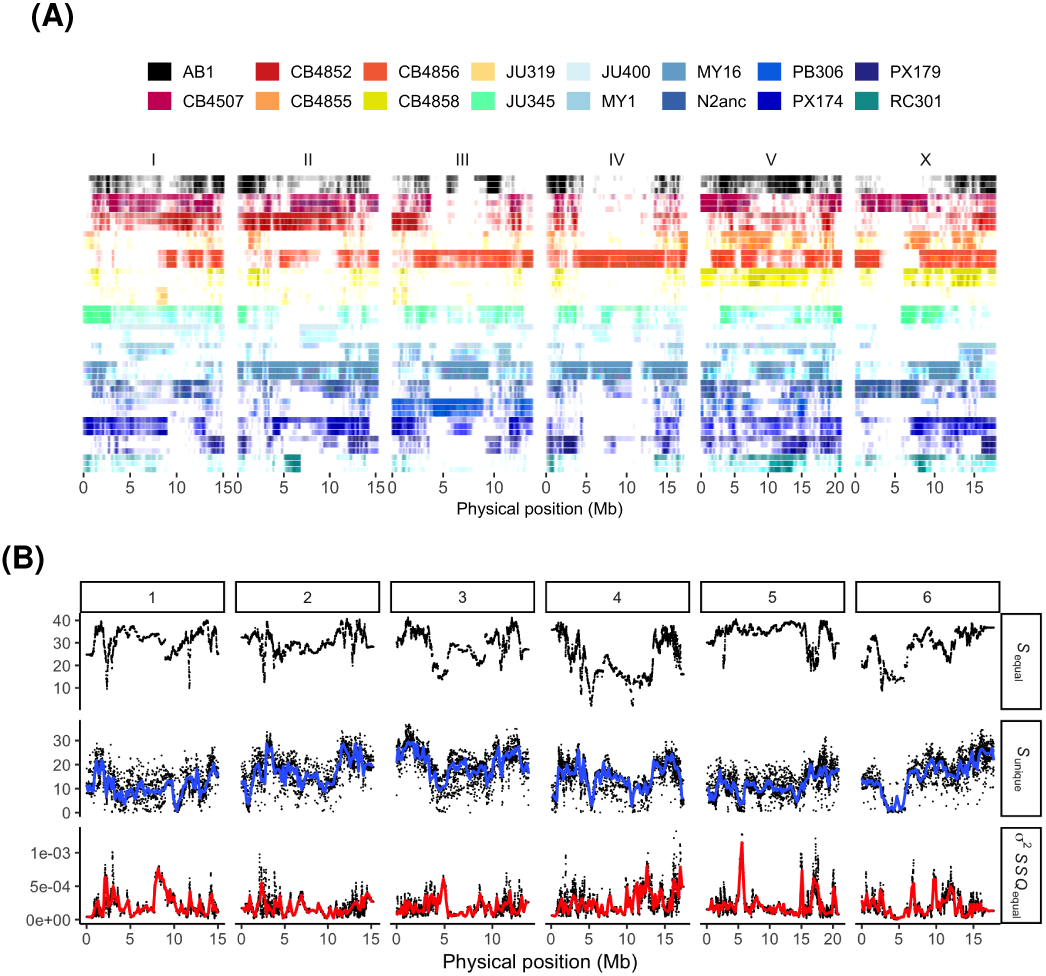
**A.** Founder representation across CeMEE populations. For each of the 16 founders, the frequency of reconstructed haplotypes is shown for A6140 (top), CA[1-3]50 (middle) and CA[1-3]100 (bottom) RILs. **B.** Upper panel: haplotype entropy across all CeMEE RILs for all 16 founders (high values indicating high diversity, or more equal founder proportions). Middle panel: as above, but for unique founder haplotypes in each 5Kb window, with a loess polynomial regression in blue. Lower panel: haplotype differentiation across population replicates (variance of the per-population sum of squared deviations from equal founder frequencies, loess fit in red. GT populations are outliers for a locus on the right tip of chromo-some V and were excluded, as were the 5 GM RILs).

### Population structure

Principal components analysis was carried out on additive genetic relatedness matrices (**A**) constructed from segregating SNPs (marker set 1 data, filtered to sites with a maximum of 2% heterozygous calls in marker set 2) with the base R function *prcomp* (R Core Team 2018). Given the matrix of *n* lines by *m* SNPs, genetic similarity was calculated by scaling each marker to mean 0 and variance 1, with **A** = **XX**^*T*^/*m*. Populations were decomposed jointly across chromosomes or recombination rate domains (Rockman and Kruglyak 2009), excluding the 5 GM RILs.

### QTL mapping simulations

#### Additive tests

Additive QTL were simulated varying heritability of a single focal marker and a background polygenic component (spread over 100, 500 or 1000 markers) with total heritability set to 0.5, and effect sizes drawn from the standard normal distribution. We fit set-based linear mixed effects models (LMMs) with random effects for additive genetic relatedness in a focal window (*R*_*w*_) and genome-wide (*R*_*g*_), across 5Kb non-overlapping windows with a minimum of 3 variants per window. The set test captures potential haplotype-based causal effects and also scales to multiple traits (Casale *et al.* 2015). In the single trait case, with mean-centered phenotype vector *y*, fixed effects *X* (here an intercept only) and residual *e*, we have:

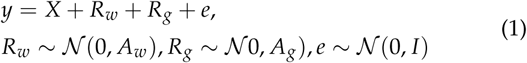

Markers were pruned of complete LD within chromosomes (*r*^2^ < 0.99) to generate the realized genetic relationship matrix *Ag* and before simulation markers were further pruned to (*r*^2^ < 0.98) for computational efficiency. For each heritability scenario (defined by the focal marker *h*^2^ and the number of markers in the polygenic component) we ran 1000 simulations sampling loci at random and recording the likelihood ratio (LR) for the full model to a null model omitting *R*_*w*_. P-values were calculated from null permutations of genotypes within *R* (30 for each test), pooling LRs across all tests for a heritability scenario. Test statistics are χ ^*2*^ distributed with a mixture of 0 and >0 degrees of freedom (Self and Liang 1987). This mixture was estimated from null LRs, then used to converting alternate LRs to p-values based on the null distribution (Listgarten *et al.* 2013; Casale *et al.* 2015). Genome-wide significance was set from the empirical QTL false discovery rate (FDR), with a QTL defined as the region of contiguous windows within -log10(p)-1.5 of the peak window. A threshold of 5 × 10^−4^ was sufficient to control the genome-wide FDR below 1% for simulated marker heritabilities of up to 0.15.

Models were implemented in python 2.7.12 with limix 1.0.12 (https://github.com/PMBio/limix).

#### Interaction test

The power to map interactions was tested by sampling marker pairs and simulating phenotypes with a given epistatic heritability. Main effects and polygenic heritability were not simulated. Markers were first pruned of strong local LD (marker set 1 with *r*^*2*^ < 0.5, MAF > 5%), and sampled pairs were tested only if all four genotype classes were present in at least 3 lines. With mean-centered phenotype vector *y*, a full model *M*1 was tested against the null additive model *M*0 by likelihood ratio test.

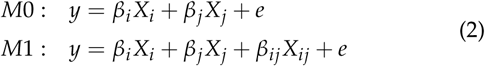

Empirical p-values were obtained by sampling responses from the null additive model (Bůžková *et al.* 2011). For each heritability level, 1000 simulations were run, taking 100 null samples at each test and then pooling these to estimate p-values. Genome-wide significance was then declared at 10% FDR after Bonferroni correction for multiple tests based on the effective number of markers (*M*_*eff*_), the number of eigenvalues of the marker correlation matrix *R* that explain at least 99% of the variance in linkage disequilibrium (as in Davis *et al.* (2016)). Covariance matrices sampled from an unobserved population are often poorly behaved and biased in the distribution of eigen-values when the number of markers (or phenotypes) *p* is much larger than the number of observations *n*, as is common using genomic data (Meyer 2016). Following Davis *et al.* (2016), we applied the non-parametric Ledoit-Wolf shrinkage estimator to *R* before eigenvalue decomposition.

## Results and discussion

### Panel composition

Through derivation of inbred lines from new experimentally evolved populations and (re)sequencing of RILs from existing populations, the CeMEE panel was expanded by almost 50% over the version 1 release (Noble *et al.* 2017). A total of 733 lines passed quality control on sequencing depth, zygosity (whether due to residual heterozygosity, or line or DNA contamination during inbreeding and sequencing), genomic relatedness, and haplotype reconstruction likelihood (Table 1).

### Improved QTL mapping power and resolution

The aims of a long-term evolution experiment and those of QTL mapping are not fully aligned. Drift and selection acting on standing genetic variation will lead to divergence among populations, and potentially to loss of genetic diversity. We showed previously extensive divergence from the founders during the initial multiparent funnel and domestication phases, with more than 32,000 SNPs fixing. Few large-scale hard sweeps were observed, however, with the loss of founder singletons accounting for around 80% of these cases, and over 97% of the autosomal genome remaining genetically variable (at 20Kb scale). Although the new CA100 populations are slightly more homogeneous than their CA50 progenitors (Figure 2A), suggesting reduced *Ne,* we continue to see 0 fixed SNP differences between any pair of replicate populations.

Mapping resolution is limited by effective recombination. With the addition of up to 3200 meioses per autosome from the 128 CA[1-3]100 RILs alone, we expect gains in both power and resolution, subject to the maintenance of recombinant diversity within and among population replicates. Potential gain (or loss) of power due to atomization of linked quantitative trait nucleotides (QTN) of antagonistic (or similar) effect is an empirical question (Bernstein *et al.* 2018). Realized genetic map expansion *(Ra)* estimated during joint haplotype reconstruction shows continued gains in the CA[1-3]100 (Figure 2B). These are seen for all chromosomes (Figure 2C), and are of a similar magnitude to the preceding 50 generations of adaptation from the A6140 in the CA[1-3]50 populations. The per-generation and chromosome increase in *Ra* is 1.21 for CA50s versus 1.15 for CA100s, with progressive underestimation of the true map expansion expected with increasingly fine recombination (Noble *et al.* 2017).

Simulations of additive QTL show, as expected, improved power over CeMEE v1, reaching 80% power for variants explaining around 3.5% of the phenotypic variance. For the set-based test with non-overlapping windows, the median detected QTL interval is 5-10Kb (1-2 windows) on chromosome arms and 10-20Kb on the centers. In general, given the relatively flat minor allele frequency (MAF) spectrum, frequency has very little effect on power to map additive QTL beyond MAF 5%.

Detecting epistasis is difficult due to scaling of the number of tests and the joint allele frequency spectrum (Phillips 2008). Only cases of second-order epistasis with very strong associations, such as those mapped previously for fitness and hermaphrodite body size in CeMEE v1 (Noble *et al.* 2017), are likely to be detected with any confidence (>50% power for interactions explaining 10% of trait variance).

### Population structure

The structured nature of the panel presents some challenges for mapping the causal basis of trait variance, particularly for redundant genetic architectures of highly polygenic traits drifting within replicate experimental lineages. For less polygenic traits, the influence of phenotype and genotype confounding on false positives weakens as the number of discrete populations increases (Rosenberg and Nordborg 2006). Structure due to experimental evolution is, of course, known by design, and in the simplest case can be handled by conditioning on population means (on the assumption of consistent directional effects across populations). Subtler patterns of realized genetic relatedness, approximated by genome-wide SNP data, can be accounted for in the standard linear mixed-effects model framework.

We previously showed that while the experimental design has, as expected, generated significant structure (Chelo and Teotónio 2013; Noble *et al.* 2017), most of the major axes of variation are not obviously associated (with the exception of GT populations, which show strong differentiation for an introgressed sex-determining allele on chromosome V captured by the first principal component). Revisiting this with new data, we again see that the principal components of additive genomic relatedness show strong structure that varies within and across chromosomes, but is largely consistent across populations (Figure 4). Structure is not significantly associated with traits such as body size and fertility (Noble *et al.* 2017), and thus may represent convergent phenotypic evolution from polygenic, yet discontinuous, genetic architectures.

Haplotype divergence varies markedly across and within chromosomes Figure 5. This is due in part to recombination rate variation among chromosomes, existing population genetic structure in the wild founders (Andersen *et al.* 2012), and selection during experimental evolution. We noted fixation of N2-like haplotypes across a large region of chromosome X centred on *npr-1* (4.77 Mb) and near-fixation of the JU345 haplotype over the *zeel-1/peel-1* selfish genetic element on chromosome I (2.35 Mb), but similar levels of divergence are seen at several uncharacterized loci across the autosomes Figure 5B, including several regions that are not conserved in the genus and appear to harbour within-species copy number variation.

## Conclusions

We have expanded the *C. elegans* multiparental experimental evolution (CeMEE) panel by almost 50%, with recombinant inbred lines drawn from discrete populations separated for 50-150 generations since common ancestry. Despite the highly structured nature of the panel, allele frequency differentiation among populations is limited, and many of the dominant axes of variation stem from genetic structure already present in the ancestral A6140 population, maintained through hundreds of generations under varying evolutionary regimes.

QTL mapping power and resolution will continue to improve as additional RILs are sequenced, though returns diminish, and mapping efforts for very highly polygenic traits may be better served by focusing on a single source population. Another possible venue for improvement is integration with the *C. elegans* natural diversity (CeNDR) panel, a growing collection of wild isolates and associated trait data (Cook *et al.* 2017). Comparing QTL mapping results from CeMEE and CeNDR holds promise to understand the importance of dominance and epistasis in adaptive responses (Barton 2017), particularly the role of genetic incompatibilities Seidel *et al.* (2008), as the former suffers from inbreeding depression (Chelo *et al.* 2014), while the latter shows outbreeding depression (Dolgin *et al.* 2007). The panel’s greatest utility for understanding trait genetics and evolution may be realised as molecular, cellular and organismal phenotypes are generated and analysed jointly (Houle *et al.* 2010).

## Acknowledgements

We thank A. Crist and V. Pereira for support with worm handling, sample preparation, and data acquisition; members of the Teotónio, Rockman and Félix labs for helpful discussion, and E. Andersen for discussion and sharing of sequencing data.

## Author contributions

RIL derivation and sequencing: L.N., M.R., H.T.; data analysis: L.N.; manuscript: L.N., M.R., H.T.

